# p53 secures the normal behavior of H3.1 histone in the nucleus by regulating nuclear phosphatidic acid and EZH2 during the G1/S phase

**DOI:** 10.1101/2023.06.27.545208

**Authors:** Tsukasa Oikawa, Junya Hasegawa, Haruka Handa, Naomi Ohnishi, Yasuhito Onodera, Ari Hashimoto, Junko Sasaki, Takehiko Sasaki, Koji Ueda, Hisataka Sabe

## Abstract

Histones are key molecules of epigenetic regulation and inheritance, and are thought to be chaperoned and transported into the nucleus appropriately prior to being integrated into nucleosomes. H3.1 histone is predominantly synthesized and enters the nucleus during the G1/S phase of the cell cycle, as a new component of duplicating nucleosomes. Here we found that p53 is necessary to secure the normal behavior and modification of H3.1 in the nucleus during the G1/S phase, in which p53 increases C-terminal domain nuclear envelope phosphatase 1 (CTDNEP1) levels and decreases enhancer of zeste homolog 2 (EZH2) levels in the H3.1 interactome. In the absence of p53, H3.1 molecules tended to be tethered at or near the nuclear envelope (NE), where they were predominantly trimethylated at lysine 27 (H3K27me3) by EZH2, without forming nucleosomes. This accumulation was likely caused by the high affinity of H3.1 towards phosphatidic acid (PA). p53 reduced nuclear PA levels by increasing levels of CTDNEP1, which activates lipin to convert PA into diacylglycerol. Induction of the *TMEM255A* gene by p53 linked p53 with CTDNEP1, in which TMEM255A stabilized CTDNEP1. We moreover found that the cytosolic H3 chaperone HSC70 attenuates the H3.1-PA interaction, and our molecular imaging analyses suggested that H3.1 molecules may be anchored around the NE after their nuclear entry. Our results expand our knowledge of p53 function in regulation of the nuclear behavior of H3.1 during the G1/S phase, in which p53 may primarily target nuclear PA and EZH2.

## Introduction

Molecular processes involved in the nuclear transport and intranuclear behavior of histones, and those involved in the engagement of histones with the nucleosome and their epigenetic marking are thought to be closely associated with genome regulation and integrity. From this perspective, molecular chaperones and transporters involved in the nuclear transport of H3-histones have been extensively studied (Alvarez et al., 2011; Apta-Smith et al., 2018; Campos et al., 2010; Campos et al., 2015; Groth et al., 2005; Mühlhäusser et al., 2001; Pardal and Bowman, 2022; Soniat et al., 2016; Tyler et al., 1999). H3.1 histone is predominantly synthesized and enters the nucleus during the G1/S phase of the cell cycle (Mendiratta et al., 2019). Histone chaperone networks, including HSC70 and chromatin assembly factor 1 (CAF1), have been shown to guide H3.1 from the cytosol to sites of DNA replication (Campos et al., 2010; Pardal and Bowman, 2022; Tagami et al., 2004). However, there are still many missing links in our understanding of the nuclear regulation and behavior of histones.

To understand the nuclear regulation and behavior of histones, we here focused on H3.1, because of its specificity to the G1/S phase of the cell cycle, which may be able to simplify experimental settings compared with the analysis of other histones. We found that p53, a product of the tumor suppressor gene *TP53*, is integral for ensuring the normal behavior of H3.1 in the nucleus, whereas p53 does not appear to be necessary for the nuclear entry of H3.1. Our results demonstrate a novel association between p53 and H3.1, in which p53 is required for H3.1 not to be trapped around the nuclear envelope (NE), and not to be therein marked by enhancer of zeste homolog 2 (EZH2) as suppressive, without forming nucleosomes. For this function, p53 appears to downregulate nuclear phosphatidic acid (PA) levels, likely via its transcriptional activity, and exclude EZH2 from the H3.1 interactome.

## Results

### Loss of p53 causes the perinuclear accumulation of lysine 27-trimethylated histone H3 (H3K27me3)

Using super-resolution three-dimensional structured illumination microscopy (SIM), we found that silencing of *TP53* by its siRNAs (si*TP53*) in normal human mammary epithelial HMLE cells causes the accumulation of H3K27me3, but not lysine 4-trimethylated histone H3 (H3K4me3) or lysine 27-acetylated histone H3 (H3K27ac) near the NE (Figure 1A, B and C, Figure 1–—figure supplement 1A). H3K27me3, H3K4me3, and H3K27ac were spread throughout the nucleus in control cells (*i.e.*, si*Scr*-treated cells) (Figure 1A). si*TP53* treatment of the human cancer cell lines A549 and MCF7 also caused similar perinuclear accumulation of H3K27me3 (Figure 1C, Figure 1–—figure supplement 1A and B). Levels of these histone modifications were not notably affected by si*TP53* (Figure 1D). Similar accumulation of H3K27me3 was also observed in the human cancer cell line H1299, which lacks p53 expression, and in the mouse embryonic fibroblast cell line MB352 established from *Trp53*^−/−^ mice (Figure 1E, Figure 1–—figure supplement 1A). We confirmed that the introduction of wild-type p53 (p53 WT) in MB352 cells cancels the H3K27me3 accumulation (Figure 1–—figure supplement 1A, C, and D). Therefore, p53 deficiency may cause the aberrant perinuclear accumulation of H3K27me3 in different types of cells, including cancer cells, in humans and mice.

**Figure 1.**
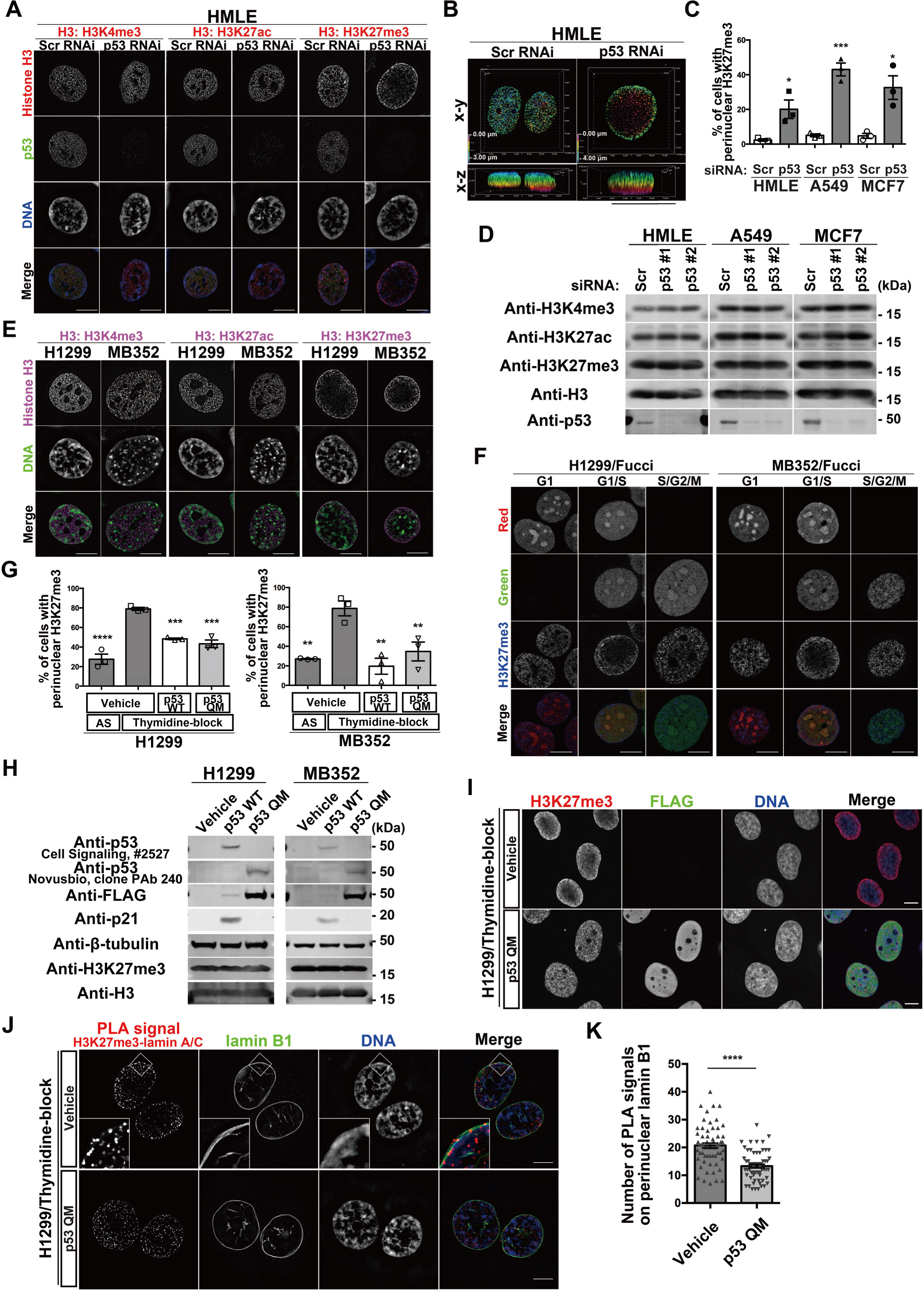
p53 suppresses perinuclear H3K27me3 during the G1/S phase. **A**, Representative SIM images of HMLE cells showing the localization of H3K4me3, H3K27ac, or H3K27me3 (red), p53 (green), and DNA (blue). Bars, 10 µm. **B**, Depth-coded alpha-blending views of H3K27me3 from the 3D-SIM images of HMLE cells. Bar, 10 µm. **C**, Quantification of perinuclear H3K27me3 accumulation in the indicated cells. n = 3 biological replicates, ≥ 100 cells per replicate; \*\*\**P* = 0.0006, \**P* < 0.03; two-tailed, unpaired *t*-test. **D**, Representative immunoblots of the indicated antibodies. **E**, Representative SIM images showing the localization of H3K4me3, H3K27ac, or H3K27me3 (magenta) and DNA (green) in the indicated cells. Bars, 10 µm. **F**, Representative SIM images showing the localization of H3K27me3 (blue) in different stages of the cell cycle, in H1299 cells and MB352 cells. Bars, 10 µm. **G**, Quantification of the indicated cells with perinuclear H3K27me3 accumulation. AS, asynchronous. n = 3 biological replicates, ≥ 50 cells per replicate; \*\*\*\**P* < 0.0001, \*\*\**P* < 0.001, and \*\**P* < 0.01; one-way ANOVA followed by Dunnett’s multiple comparisons test. **H**, Representative immunoblots of the indicated antibodies. **I**, Representative confocal images of H1299 cells showing the localization of H3K27me3 (red), FLAG (green), and DNA (blue) Bars, 10 µm. **J**, Representative SIM images of H1299 cells showing the localization of PLA signals between H3K27me3 and lamin A/C (red), lamin B1 (green), and DNA (blue) Bars, 10 µm. **K**, Quantification of the PLA spots that overlap with perinuclear lamin B1 in each nucleus. n = 3 biological replicates, ≥ 60 nuclei; \*\*\*\**P* < 0.0001; two-tailed, unpaired *t*-test.

### Perinuclear accumulation of H3K27me3 occurs during the G1/S phase

We then investigated whether H3K27me3 accumulation occurs during a specific phase of the cell cycle. H1299 cells and MB352 cells stably expressing the cell cycle indicator Fucci (Sakaue-Sawano et al., 2008) demonstrated that H3K27me3 accumulation occurs predominantly from the late G1 to the early S phase in the absence of p53 (Figure 1F, Figure 1–—figure supplement 1A). Thymidine blockade of G1/S progression in H1299 and MB352 cells also caused H3K27me3 accumulation (Figure 1G, Figure 1–—figure supplement 1A and E). Therefore, the perinuclear accumulation of H3K27me3 in the absence of p53 appeared to occur predominantly during the G1/S phase.

### p53 suppresses H3K27me3 accumulation without cell-cycle arrest or apoptosis

The expression of p53 WT significantly decreased H3K27me3 accumulation in thymidine-blocked H1299 cells and MB352 cells (Figure 1G, Figure 1–—figure supplement 1A and D). However, p53 is known to induce genes involved in cell-cycle arrest or apoptosis (el-Deiry et al., 1993) (Figure 1H, Figure 1–—figure supplement 1E). We then used a p53 mutant, p53 QM, bearing the L22Q/W23S/W53Q/F54S mutations, which was previously shown to not induce the expression of such genes (see Figure 1–—figure supplement 1D) (Venot et al., 1999). We here confirmed that p53 QM did not induce genes known to be involved in cell-cycle arrest, apoptosis, and metabolism, when expressed in H1299 cells and MB352 cells (Figure 1H, Figure 1–—figure supplement 1D and E, Figure 1–—figure supplement 2, Supplementary file 1). Nevertheless, p53 QM substantially suppressed the accumulation of H3K27me3 in these cells (Figure 1G and I, Figure 1–—figure supplement 1A), whereas it did not affect the localization of another NE-associated histone, lysine 9-trimethylated histone H3 (H3K9me3) (Figure 1–—figure supplement 1F). Furthermore, the proximity ligation assay (PLA) coupled with SIM imaging in thymidine-blocked H1299 cells demonstrated that the expression of p53 QM significantly reduces the colocalization of H3K27me3 with lamin A/C near the NE (Figure 1J and K). Therefore, p53 appears to suppress the aberrant perinuclear accumulation of H3K27me3 and its colocalization with lamin A/C during the G1/S phase, likely independent of cell-cycle arrest, apoptosis, and metabolic remodeling.

### Properties of perinuclearly accumulated H3K27me3 histones

We then investigated the properties of the perinuclearly accumulated H3K27me3 histones. Costaining of H3K27me3 with lamin A/C in H1299 cells showed their close colocalization with the NE (Figure 2A). Their costaining with DNA suggested that these histones do not colocalize well with DNA (Figure 2B). We then visualized the DNA replication sites using biotin-labeled deoxyuridine triphosphate (dUTP) (Alabert et al., 2014; Maya-Mendoza et al., 2012). SIM imaging of H1299 cells, which were blocked once with thymidine and then released for 20 min, revealed that most of the perinuclearly accumulated H3K27me3 histones were not localized to DNA replication sites, whereas the localization of H3K27me3 histones to DNA replication sites was clearly detected in the presence of p53 QM (Figure 2C and D).

**Figure 2.**
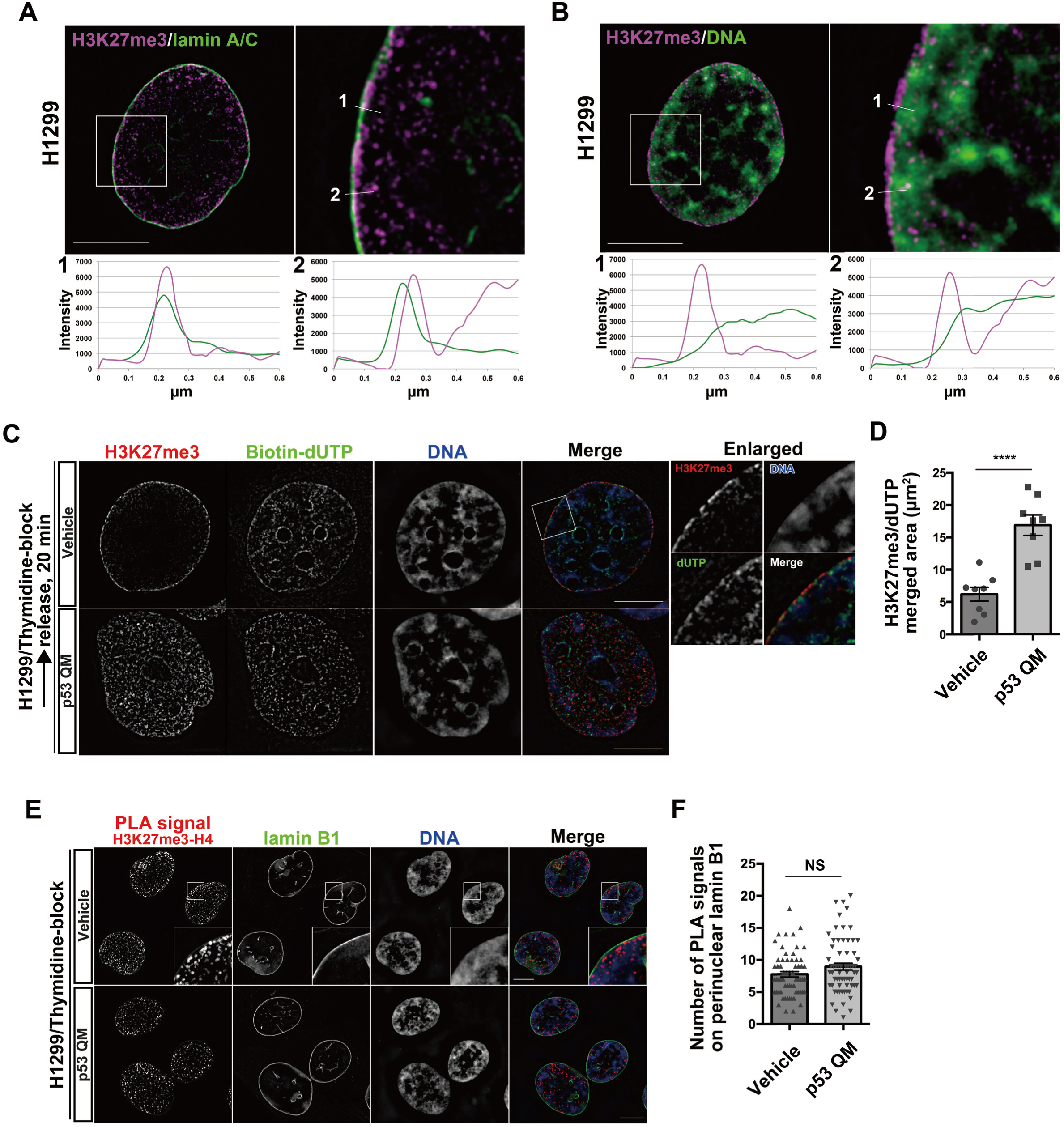
Perinuclear H3K27me3 does not form nucleosomes. **A**, **B**, Representative SIM images of a H1299 cell showing the localization of H3K27me3 (magenta) and lamin A/C (green) (**A**), or H3K27me3 (magenta) and DNA (green) (**B**). Enlarged images of the white boxed areas are shown on the right. Pixel intensities on the traversing white lines (1 and 2) are shown at the bottom. Bars, 10 µm. **C**, Representative SIM images of H1299 cells showing the localization of H3K27me3 (red), biotin-dUTP (green), and DNA (blue). Enlarged images of the white boxed area are shown on the right. Bars, 10 µm. **D**, Quantification of the area of H3K27me3 and dUTP colocalization. n = 2 biological replicates; \*\*\*\**P* < 0.0001; two-tailed, unpaired *t*-test. **E**, Representative SIM images of H1299 cells showing the localization of the PLA signals between H3K27me3 and H4 (red), lamin B1 (green), and DNA (blue). Bars, 10 µm. **F**, Quantification of the PLA spots that overlap with perinuclear lamin B1 in each nucleus. n = 3 biological replicates; ≥ 60 nuclei; NS, not significant; two-tailed, unpaired *t*-test

We furthermore analyzed the possible nucleosome formation of these histones. PLA between H3K27me3 and H4 in thymidine-blocked H1299 cells demonstrated that most of the PLA signals were found inside the nucleoplasm, rather than near the NE (Figure 2E and F). Furthermore, the levels of these signals near the NE were not notably changed, irrespective of p53 QM (Figure 2E and F). Thus, most of the H3K27me3 histones accumulated near the NE in the absence of p53 did not appear to have formed nucleosomes, bound to DNA, or localized to DNA replication sites.

### H3.1/H3.2 comprises the accumulated H3K27me3

H3.1 and H3.2 are synthesized in the cytosol, and then transported into the nucleus during the G1/S phase of the cell cycle, whereas H3.3 acts independently of DNA replication (Mendiratta et al., 2019; Tagami et al., 2004). These H3 variants were expressed at similar levels in H1299 cells, irrespective of p53 QM expression or thymidine blockade (Figure 3A). Using antibodies specific to H3.1/H3.2 and to H3.3, we then found that H3.1/H3.2, but not H3.3, accumulated near the NE in thymidine-blocked H1299 cells (Figure 3B), and that such accumulation was largely mitigated by p53 QM (Figure 3C). PLA coupled with SIM imaging in thymidine-blocked H1299 cells demonstrated that the expression of p53 QM significantly reduces the colocalization of H3.1/H3.2 with lamin A/C near the NE (Figure 3D and E). Thus, H3.1/H3.2, but not H3.3, appear to predominantly comprise the perinuclearly accumulated H3K27me3 histones. Antibodies that selectively recognize either H3.1 or H3.2 were not available, and it hence remains unclear as to which of them comprise the accumulated H3K27me3 histones.

**Figure 3.**
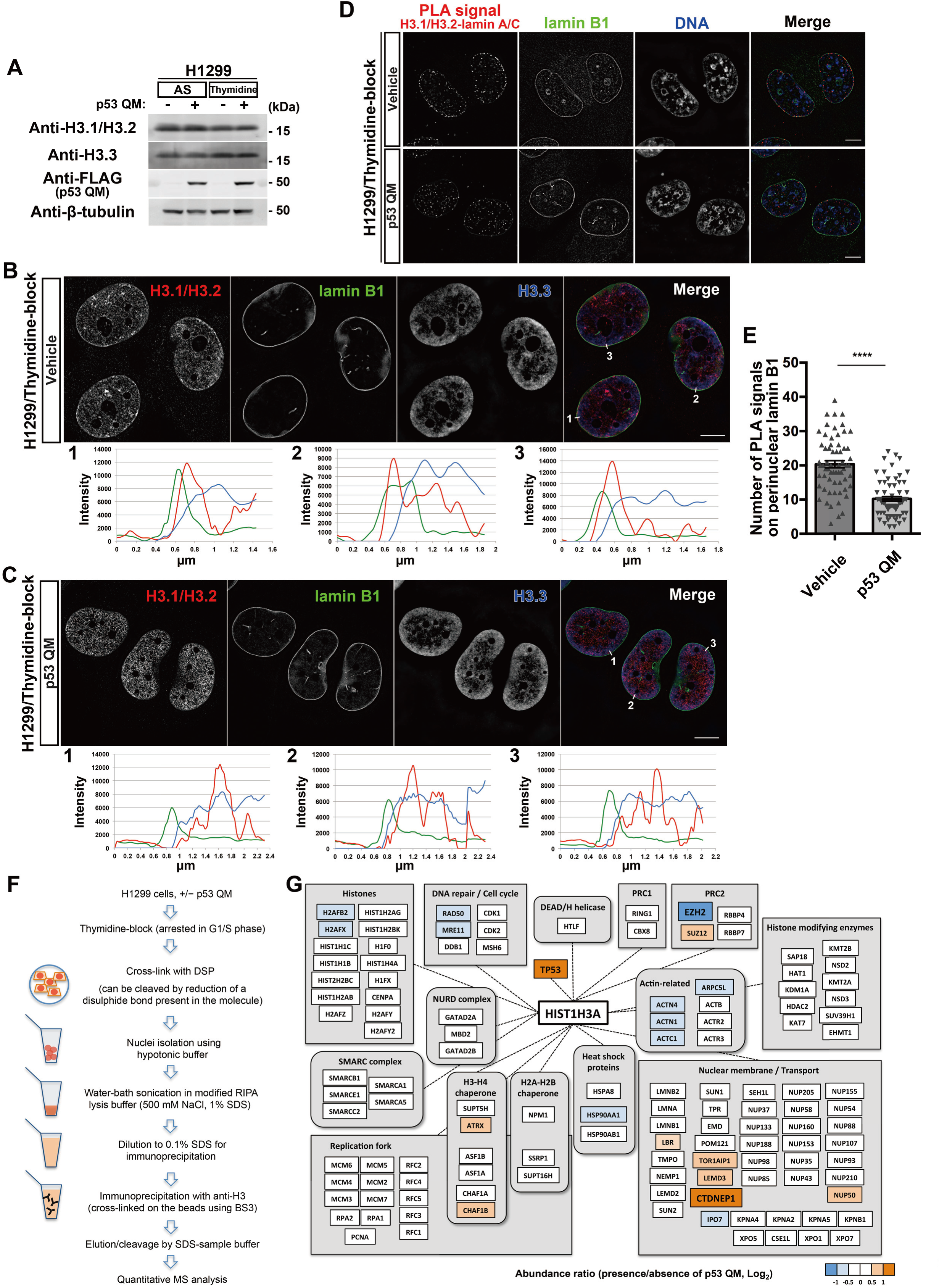
H3.1/H3.2 comprises perinuclearly accumulated H3K27me3. **A**, Representative immunoblots of the indicated antibodies. AS, asynchronous. **B**, **C**, Representative SIM images of H1299 cells showing the localization of H3.1/H3.2 (red), lamin B1 (green), and H3.3 (blue) in the absence (**B**) or presence (**C**) of p53 QM. Pixel intensities on the traversing white lines (1, 2, and 3) are shown at the bottom. Bars, 10 µm. **D**, Representative SIM images of H1299 cells showing the localization of the PLA signals between H3.1/H3.2 and lamin A/C (red), lamin B1 (green), and DNA (blue). Bars, 10 µm. **E**, Quantification of the PLA spots that overlap with perinuclear lamin B1 in each nucleus. n = 3 biological replicates; ≥ 60 nuclei; \*\*\*\**P* < 0.0001; two-tailed, unpaired *t*-test. **F**, Schematic representation of H3 immunoprecipitation followed by quantitative MS analysis. **G**, Representative proteins identified from the MS analysis. The relative abundance of the proteins is presented as a ratio (Log_2_) in the presence/absence of p53 QM.

### p53 increases CTDNEP1 and decreases EZH2 levels in the nuclear H3.1 interactome

We then sought to obtain a clue to link p53 with H3.1/H3.2. For this purpose, we performed an interactome analysis using liquid chromatography-mass spectrometry (LC-MS) on anti-H3 immunoprecipitants from the nuclear fraction of thymidine-blocked H1299 cells (Figure 3F). Proteins were crosslinked with dithiobis(succinimidyl propionate) (DSP) before solubilization. Notably, we found that the anti-H3 antibody predominantly pulled down H3.1 under this condition, rather than H3.2 or H3.3 (Figure 3G, Supplementary file 2), although we do not have a clear explanation for this. Among the different proteins that were coprecipitated, the amount of C-terminal domain nuclear envelope phosphatase 1 (CTDNEP1) was more than 4-fold higher in the presence of p53 QM than in its absence (Figure 3G, Supplementary file 2). EZH2 was also coprecipitated, but its amount was about 2-fold lower in the presence of p53 than in its absence (Figure 3G, Supplementary file 2). The amount of H3.1 in the immunoprecipitants was not notably changed irrespective of the presence/absence of p53 QM; and p53, when present, was coprecipitated with H3.1 (Figure 3G, Supplementary file 2). Thus, p53 may play a role in increasing CTDNEP1 and decreasing EZH2 levels in the nuclear H3.1 interactome formed during the G1/S phase.

### p53 suppresses the perinuclear interaction of H3.1 with EZH2

We then focused on EZH2. EZH2 is the catalytic subunit of polycomb repressive complex 2, which generates H3K27me3 (Cao et al., 2002). We confirmed the above results by western blotting, in which a larger amount of EZH2 was detected in anti-H3 immunoprecipitants from the G1/S-arrested cells in the absence of p53 QM (Figure 4A). Benzonase is a potent endonuclease, and benzonase treatment during immunoprecipitation did not affect the amount of coprecipitated EZH2, avoiding the possible contamination of nuclear DNA (Figure 4A).

**Figure 4.**
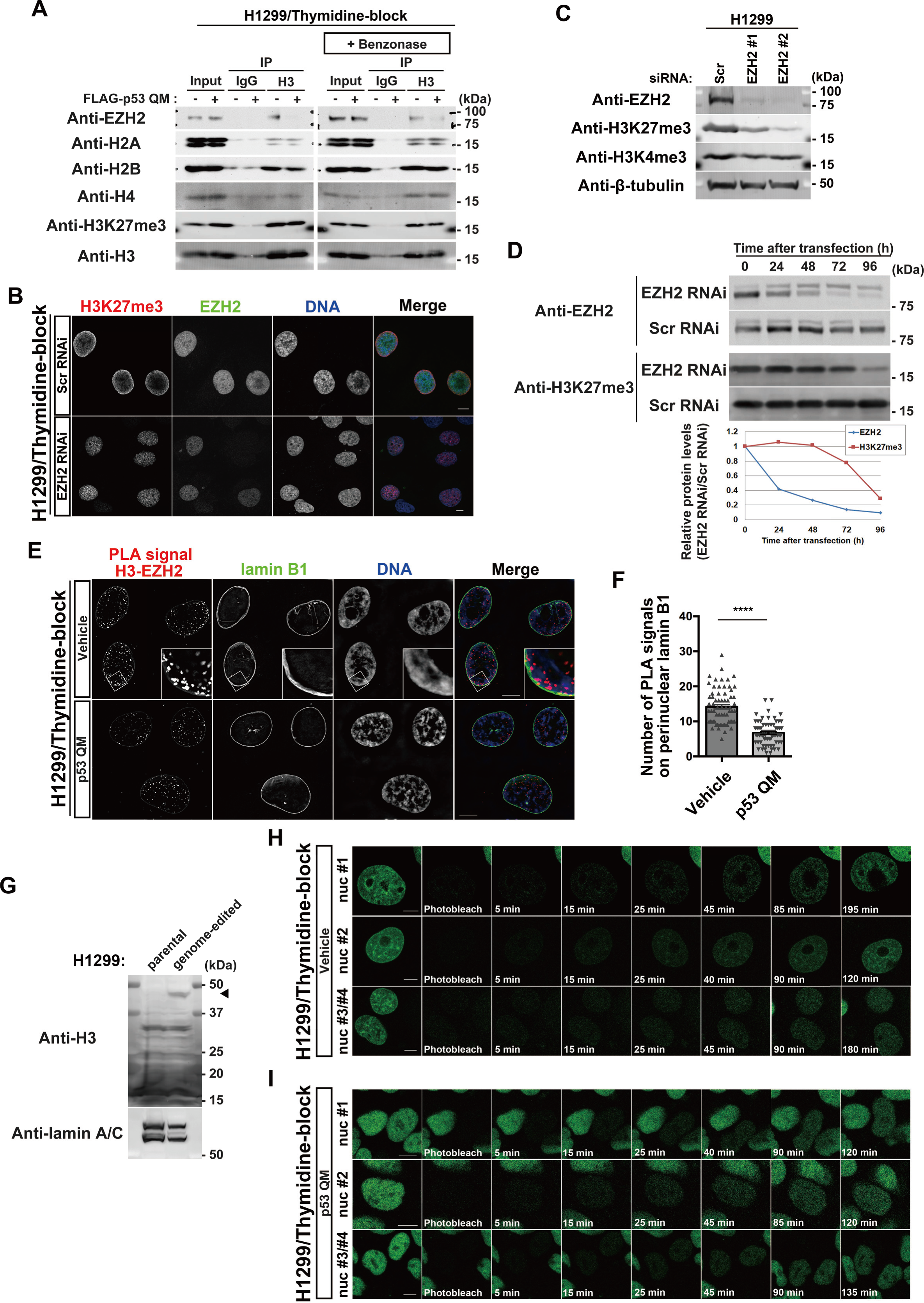
p53 reduces the proximity between EZH2 and perinuclearly tethered H3. **A**, Representative immunoblots of the indicated antibodies. IP, immunoprecipitation. **B**, Representative confocal images of H1299 cells showing the localization of H3K27me3 (red), EZH2 (green), and DNA (blue). Bars, 10 µm. **C**, Representative immunoblots of the indicated antibodies. **D**, Representative immunoblots of the indicated antibodies. Normalized relative protein levels (EZH2 RNAi/Scr RNAi) are shown at the bottom. **E**, Representative SIM images of H1299 cells showing the localization of PLA signals between H3 and EZH2 (red), lamin B1 (green), and DNA (blue). Bars, 10 µm. **F**, Quantification of the PLA spots that overlap with perinuclear lamin B1 in each nucleus. n = 3 biological replicates, ≥ 60 nuclei; \*\*\*\**P* < 0.0001; two-tailed, unpaired *t*-test. **G**, Representative immunoblots of the indicated antibodies. The arrowhead indicates H3.1-Dronpa. **H**, **I**, Representative confocal images showing the localization of H3.1-Dronpa in the absence (**H**) or presence (**I**) of p53 QM in H1299 cells. Bars, 10 µm.

si*EZH2* substantially mitigated the perinuclear accumulation of H3K27me3 in these thymidine-blocked H1299 cells (Figure 4B). Western blot analysis showed that si*EZH2* reduced the total cellular amounts of H3K27me3, but not H3K4me3 (Figure 4C). However, a small amount of H3K27me3 was still observed in some cells after si*EZH2* treatment (Figure 4B). This may be owing to a delayed decrease in H3K27me3 after *EZH2* silencing (Figure 4D). In addition, the PLA showed that p53 QM significantly suppresses the colocalization between EZH2 and H3 near the NE in G1/S-arrested cells (Figure 4E and F). Therefore, taken together, it is likely that EZH2 is responsible for the perinuclear accumulation of H3K27me3 whereas p53 appears to suppress the perinuclear interaction between H3 and EZH2.

### H3.1 molecules are tethered around the NE after their nuclear entry

There are two possibilities for the mechanism underlying the accumulation of such H3K27me3 histones that are mostly composed of H3.1/H3.2 around the NE; i.e., they were either trapped by the NE when entering the nucleus during the G1/S phase, or they were tethered around the NE after entering the nucleus. To understand which possibility is more likely, we generated H1299 cells with the gene encoding Dronpa, a fluorescent protein with reversible photobleaching/photoactivation (Ando et al., 2004) at the 3′-end of the *HIST1H3A* gene (encoding H3.1), using the microhomology-mediated end joining (MMEJ)-assisted knock-in system (Sakuma et al., 2016) (Figure 4G). We then performed fluorescence recovery after photobleaching (FRAP) analysis of H3.1-Dronpa in the nucleus (see methods). After photobleaching the nucleus of a thymidine-blocked H1299 cell, we observed a gradual increase in H3.1-Dronpa signals both at the nuclear periphery and inside the nucleus in 30 to 40 min (Figure 4H, Figure 4–—video 1A, B, and C). We observed similar recovery dynamics of the H3.1-Dronpa signals after photobleaching in the presence of p53 QM, although the perinuclear accumulation of H3.1-Dronpa was not evident (Figure 4I, Figure 4–—video 2A, B, and C). Therefore, it is likely that H3.1 molecules are tethered near the NE after entering the nucleus.

### p53 utilizes CTDNEP1 to reduce nuclear PA levels

We next focused on CTDNEP1, which was increased in the H3.1 interactome by p53. CTDNEP1 converts PA to diacylglycerol (DAG) via activation of lipin PA phosphatases (Bahmanyar and Schlieker, 2020; Barger et al., 2021; Han et al., 2012; Su et al., 2014). PA is a minor component of nuclear lipids, and mostly localizes to the NE (Bahmanyar and Schlieker, 2020). We found that H3.1 has strong affinity to PA, but not to DAG (Figure 5A). Therefore, PA may cause the accumulation of H3.1 around the NE.

**Figure 5.**
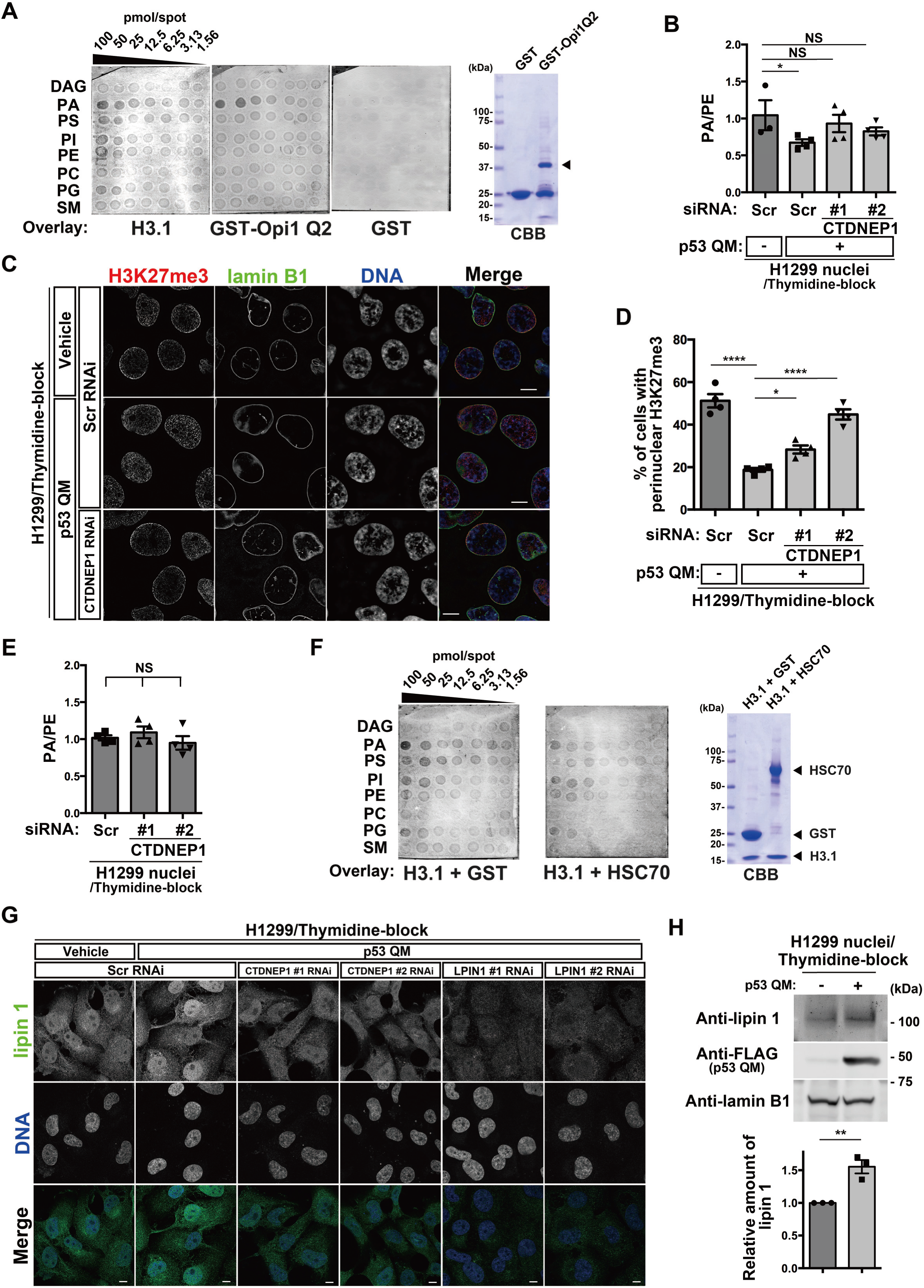
p53 and CTDNEP1 reduce nuclear PA levels by increasing lipin 1 in the nucleus. **A**, H3.1-lipid interactions clarified by membrane lipid arrays, with GST and the Q2 domain of yeast Opi1 (Opi1 Q2) as a negative and positive control, respectively. The membrane was spotted with the indicated amounts of the following lipids: diacylglycerol (DAG); phosphatidic acid (PA); phosphatidylserine (PS); phosphatidylinositol (PI); phosphatidylethanolamine (PE); phosphatidylcholine (PC); phosphatidylglycerol (PG); and sphingomyelin (SM). CBB, coomassie brilliant blue. The arrowhead indicates the band of GST-Opi1 Q2. The CBB blot on the right demonstrates the quality of the proteins used in this assay. **B**, Absolute quantification of PA and PE in the nuclei of H1299 cells. PA/PE ratios are indicated. n = 3 or 4 biological replicates; \**P* < 0.05; NS, not significant; one-way ANOVA followed by Dunnett’s multiple comparisons test. **C**, Representative SIM images of H1299 cells showing the localization of H3K27me3 (red), lamin B1 (green), and DNA (blue). Bars, 10 µm. **D**, Quantification of H1299 cells with perinuclear H3K27me3 accumulation. n = 4 biological replicates, ≥ 100 cells per replicate; \*\*\*\**P* < 0.0001, \**P* < 0.05; one-way ANOVA followed by Dunnett’s multiple comparisons test. **E**, Absolute quantification of PA and PE in the nuclei of H1299 cells. PA/PE ratios are indicated. n = 4 biological replicates; NS, not significant by one-way ANOVA. **F**, H3.1-lipid interactions in the presence of equimolar amounts of GST or HSC70 were clarified by membrane lipid arrays, as in **A**. The CBB blot on the right demonstrates the quality of the proteins used in this assay. **G**, Representative confocal images of H1299 cells showing the localization of lipin 1 (green) and DNA (blue). Bars, 10 µm. **H**, Representative immunoblots of the indicated antibodies. Normalized lipin 1/lamin B1 ratios are shown. n = 3 biological replicates; \*\**P* < 0.01; two-tailed, unpaired *t*-test.

We then tested whether CTDNEP1 reduces nuclear PA levels and the perinuclear accumulation of H3K27me3. We performed mass spectrometry analysis of nuclear PA levels using nuclear phosphatidylethanolamine (PE) as an internal control, and found that nuclear PA levels (*i.e*., the PA/PE ratio) in thymidine-blocked H1299 cells are significantly lower in the presence of p53 QM than in its absence (Figure 5B), and that si*CTDNEP1* treatment cancels this reduction (Figure 5B, Figure 5–—figure supplement 3A) and restores the perinuclear accumulation of H3K27me3 (Figure 5C and D, Figure 1–—figure supplement 1A). As a control, we confirmed that si*CTDNEP1* treatment in p53-deficient H1299 cells does not notably affect nuclear PA levels (Figure 5E, Figure 5–—figure supplement 3A).

Then, the question arises as to how H3.1 can pass through the NE when p53 is absent and PA levels increase. HSC70 (encoded by *HSPA8*) is a chaperone for H3 (Campos et al., 2010), and was included within the H3.1 interactome irrespective of the presence of p53 QM (see Figure 3G). We found that HSC70 attenuates the interaction of H3.1 with PA (Figure 5F). Therefore, HSC70 may assist H3.1 to smoothly enter the nucleus even in the absence of p53.

The nuclear levels of lipin 1 were also increased upon p53 QM expression in thymidine-blocked H1299 cells, and si*CTDNEP1* canceled this augmentation (Figure 5G, Figure 5–—figure supplement 3A). A biochemical analysis confirmed these results, in which p53 QM increased the amounts of lipin 1 in the nuclear fraction (Figure 5H). CTDNEP1 dephosphorylates and activates lipins (Han et al., 2012; Su et al., 2014). Phosphorylation levels of lipin 1 were unaffected by the presence or absence of p53 QM (Figure 5–—figure supplement 3B).

Collectively, p53 may increase CTDNEP1 levels, and CTDNEP1 may in turn increase lipin 1 and decrease nuclear PA levels, thus alleviating the abnormal accumulation of H3K27me3 near the NE. On the other hand, as antibodies against lipin 2 and lipin 3 were not available, we were unable to exclude the possible involvement of these isoforms in CTDNEP1 function.

### p53 induces *TMEM255A* to increase CTDNEP1 levels

We then sought to understand the possible molecular link between p53 and CTDNEP1. The *CTDNEP1* gene promoter does not contain p53-binding sites, and *CTDNEP1* mRNA was not increased by p53 QM in H1299 cells (Figure 6–—figure supplement 4A). We however found that si*TP53* reduces CTDNEP1 protein levels in MCF7 cells without reducing its mRNA (Figure 6–—figure supplement 4B and C). si*CTDNEP1* also reduced lipin 1 protein levels in H1299 cells (Figure 6A, Figure 5–—figure supplement 3A).

**Figure 6.**
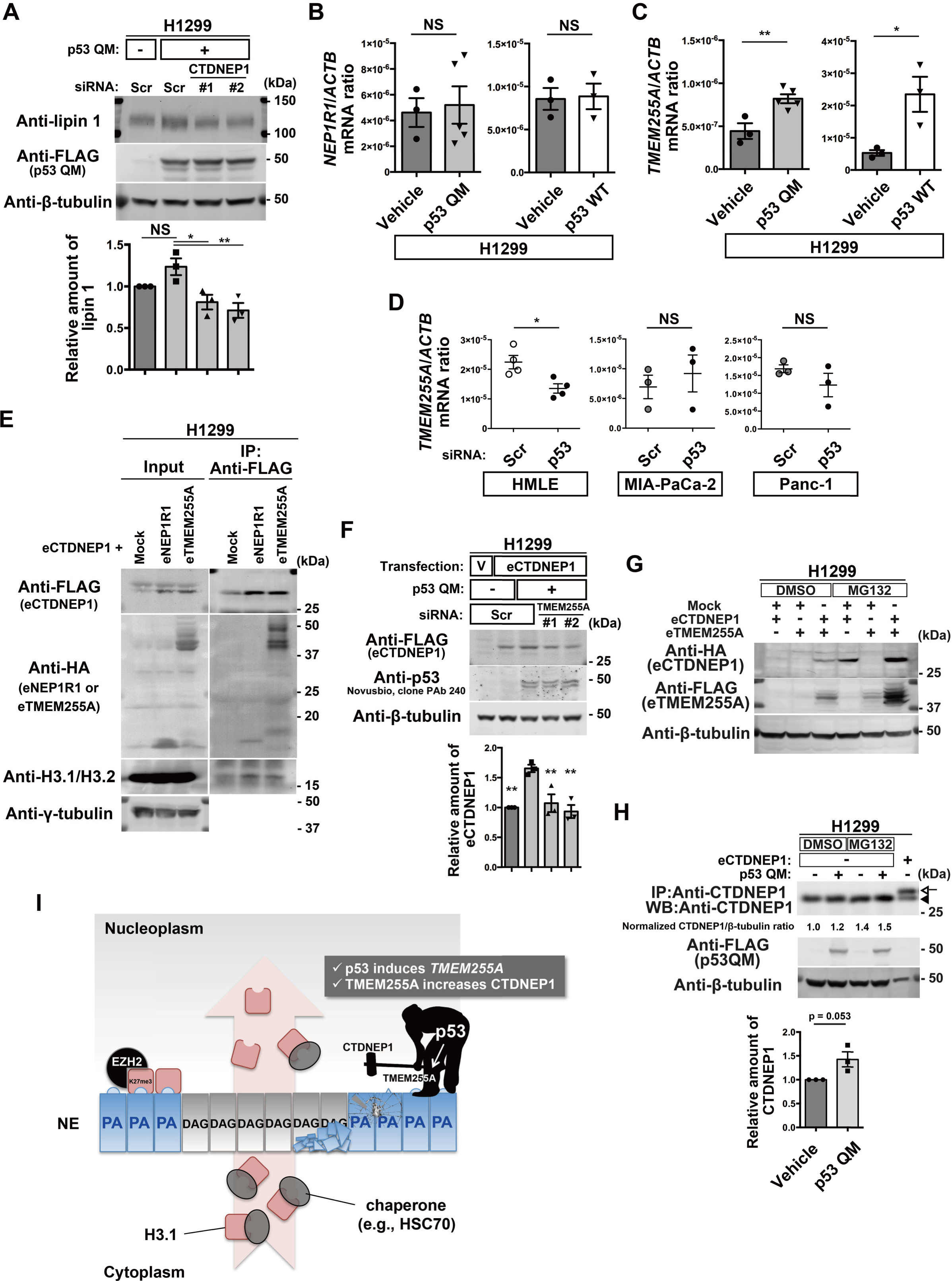
TMEM255A increases CTDNEP1 downstream of p53. **A**, Representative immunoblots using the indicated antibodies. Normalized lipin 1/β-tubulin ratios are shown. n = 3 biological replicates; \*\**P* < 0.01, \**P* < 0.05; NS, not significant; one-way ANOVA followed by Dunnett’s multiple comparisons test. **B**, **C**, Quantification of *NEP1R1* (**B**) and *TMEM255A* (**C**) mRNA (normalized by *ACTB* mRNA) in H1299 cells. n ≥ 3 biological replicates; \*\**P* < 0.01, \**P* < 0.03; NS, not significant; two-tailed, unpaired *t*-test. **D**, Quantification of *TMEM255A* mRNA (normalized by *ACTB* mRNA). n ≥ 3 biological replicates; \**P* < 0.02; NS, not significant; two-tailed, unpaired *t*-test. **E**, Representative immunoblots using the indicated antibodies. IP, immunoprecipitation. **F**, Representative immunoblots using the indicated antibodies. V, vehicle. Normalized eCTDNEP1/β-tubulin ratios are shown. n = 3 biological replicates; \*\**P* < 0.01; one-way ANOVA followed by Dunnett’s multiple comparisons test. **G**, Representative immunoblots using the indicated antibodies. **H**, Representative immunoblots using the indicated antibodies. IP, immunoprecipitation. WB, western blotting. The arrowhead indicates endogenous CTDNEP1, and the arrow indicates eCTDNEP1. Normalized CTDNEP1/β-tubulin ratios of the DMSO-treated samples are shown at the bottom. n = 3 biological replicates; *P* = 0.053; two-tailed, unpaired *t*-test. **I**, Model for the inhibition of the perinuclear accumulation of H3K27me3 regulated by p53.

CTDNEP1-regulatory subunit-1 (NEP1R1) was shown to increase CTDNEP1 protein levels by binding to CTDNEP1 (Han et al., 2012; Jacquemyn et al., 2021; Su et al., 2014). However, neither p53 QM nor p53 WT induced *NEP1R1* in H1299 cells (Figure 6B). We then found that a hitherto poorly characterized gene, namely, *TMEM255A* (also called *FAM70A*), was induced by p53 WT and p53 QM in H1299 cells (Figure 6C, Supplementary file 1). *TMEM255A* mRNA was also induced by endogenous p53 WT in HMLE cells (Figure 6D, Figure 6–—figure supplement 4D), although we have not yet clarified whether the *TMEM255A* gene is a direct target of p53. On the other hand, *TMEM255A* mRNA was not induced by mutant p53s expressed in cancer cells, such as MIA-PaCa-2 cells (expressing p53 R248W) and Panc-1 cells (expressing p53 R273H) (Figure 6D, Figure 6–—figure supplement 4D), in which the perinuclear accumulation of H3K27me3 was observed (Figure 6–—figure supplement 4E).

We then hypothesized that the TMEM255A protein increases CTDNEP1 protein levels. The expression of TMEM255A increased CTDNEP1 levels in reconstitution experiments, in which we expressed cDNAs encoding epitope-tagged TMEM255A (eTMEM255A: TMEM255A-Myc-FLAG or HA-TMEM255A) and epitope-tagged CTDNEP1 (eCTDNEP1: CTDNEP1-Myc-FLAG or HA-CTDNEP1) (Figure 6E, Figure 6–—figure supplement 4F). Moreover, eTMEM255A was clearly coprecipitated with eCTDNEP1 (Figure 6E, Figure 6–—figure supplement 4F). Furthermore, p53 QM increased eCTDNEP1 levels in H1299 cells, and si*TMEM255A* cancelled this increase (Figure 6F, Figure 6–—figure supplement 4G). Furthermore, single expression of either of these cDNAs did not lead to robust expression of the corresponding protein (Figure 6G). Therefore, CTDNEP1 appears to require TMEM255A for its protein expression. Supporting this notion, ColabFold analysis of the predicted 3D structure of protein complexes (Mirdita et al., 2022) suggested that CTDNEP1 can make a complex with TMEM255A through its transmembrane alpha helix (Figure 6–—figure supplement 4H), although we have yet to confirm this possibility biochemically. Moreover, inhibition of proteasome activity by MG132 increased protein levels of both eCTDNEP1 and eTMEM255A in H1299 cells (Figure 6G), and the level of endogenous CTDNEP1 was increased in the presence of p53 QM and MG132 (Figure 6H). Therefore, it is likely that p53 induces the *TMEM255A* gene, and its protein product may stabilize CTDNEP1 via circumventing protein degradation, possibly by their association.

On the other hand, the degree of CTDNEP1 increase by the p53-TMEM255A-CTDNEP1 axis appeared to be smaller than what we observed earlier in the H3.1 interactome. We also found substantial changes in the subcellular localization of HA-CTDNEP1; a higher amount of HA-CTDNEP1 was found to be localized around the NE in the presence of p53 QM than in its absence (Figure 6–—figure supplement 4I). Thus, mechanisms other than this might also exist by which p53 increases CTDNEP1 in the H3.1 interactome.

## Discussion

Our results described in this paper demonstrate a novel function of p53 in the regulation of nuclear lipid components, and show that this function is crucial to normalize the nuclear behavior of H3.1. We have also identified an unexpected function of EZH2 that may emerge when the above function of p53 is impaired. As a result, in the absence of p53, H3.1 molecules may be tethered around the NE and therein be marked as suppressive by EZH2 without forming nucleosomes during the G1/S phase of the cell cycle (Figure 6I).

One of the major targets of p53 in normalizing the nuclear behavior of H3.1 is the downregulation of nuclear PA levels. CTDNEP1 is primarily responsible for this downregulation, and its amounts are increased in the H3.1 interactome by p53. p53 induces *TMEM255A* gene expression, and its protein product then stabilizes the CTDNEP1 protein. Another action of p53 is the downregulation of EZH2 within the H3.1 interactome, and p53 has previously been shown to suppress the *EZH2* gene promoter (Tang et al., 2004). Thus, the duality of p53 in transcriptional regulation may underlie the herein identified function of p53. However, many issues remain unsolved. For example, the p53-TMEM255A-CTDNEP1 axis may not fully explain the p53-mediated increase in CTDNEP1 levels in the H3.1 interactome, as mentioned earlier. Total cellular EZH2 protein levels were not notably changed in H1299 cells by p53 (see Figure 4A), and hence some unknown mechanisms might exist that specifically regulate the amounts of EZH2 in the H3.1 interactome. Mechanisms by which p53 utilizes CTDNEP1 in the H3.1 interactome also await to be elucidated.

In the absence of p53, H3.1 molecules may be tethered at or be located near the NE after they have entered the nucleus, rather than be directly trapped at the NE during their entry into the nucleus. H3.1 shows robust affinity to PA, whereas its chaperone HSC70 can attenuate the H3.1-PA association. HSC70 is thought to be associated with H3.1 primarily in the cytoplasm (*i.e.*, thus originally named a “cytoplasmic” chaperone), and most nuclear H3.1 molecules were shown to be associated with other chaperones, such as CAF1 (Campos et al., 2010). Therefore, H3.1 molecules appear to be dissociated from HSC70 after entering the nucleus, and may utilize other chaperones, such as CAF1, in the nucleus. Taken together, these properties would explain to some extent why H3.1 goes through the NE and then becomes anchored around the NE in the absence of p53. Mechanisms regulating PA in the NE as well as in the nuclear pore complex have been extensively studied (Barger et al., 2021; Jacquemyn et al., 2021; Merta et al., 2021; Penfield et al., 2020; Thaller et al., 2021), because of their possible association with genetic/epigenetic control and integrity. Hence, understanding the detailed mechanisms by which PA constrains H3.1 molecules at the NE, and also the detailed mechanisms as to how EZH2 targets such H3.1 molecules being anchored to the NE without forming nucleosomes will be of particular importance. Whether such aberrantly marked H3.1 molecules will then be discarded from the nucleus, or be used in some way in the nucleosomes should also be clarified.

We have previously shown that the loss of normal p53 in epithelial cells may cause the aberrant deposition of H3K27me3 on epithelial-specific gene loci, such as *CDH1*, and that p53 utilizes EZH2 in this process (Oikawa et al., 2018a; Oikawa et al., 2018b). However, the aberrant accumulation of H3K27me3 also occurs in p53-decient mouse fibroblasts. Thus, the function of p53 in controlling the normal behavior of H3.1 does not appear to be specific to epithelial cells.

In summary, we found that there is a tight control on the nuclear behavior of H3.1 by p53, and demonstrated unprecedented perspectives regarding p53 function and EZH2-mediated H3K27me3 modification that may occur with unmethylated H3.1 molecules at or near the NE. Then, an outstanding question is, if such functions of p53 and EZH2 exist, whether any of them are associated with epigenetic control and integrity. Whether such regulations are necessary only for the G1/S-specific histones should also be clarified.

## Materials and methods

### Plasmid construction and retroviral gene transduction

A cDNA encoding a p53 protein with quadruple mutations (L22Q/W23S/W53Q/F54S; p53 QM) was generated by PCR-based mutagenesis (Hashimoto et al., 2016). The cDNA for p53 WT was cloned into pRetroX-IRES-ZsGreen1 (Clontech). The cDNA for p53 WT or p53 QM was cloned together with the DNA sequence for an NH_2_-terminal FLAG tag into pRetroX-Tight-Pur (Clontech). Retroviruses with the vesicular stomatitis virus–G (VSV-G) envelope were produced by transfection of GP2-293 cells (Clontech) with the pRetroX construct and pVSV-G (Clontech) using Lipofectamine LTX reagent (Invitrogen), following the manufacturer’s instructions. cDNAs encoding *CTDNEP1*, *NEP1R1*, and *TMEM255A* followed by the Myc-FLAG tag were purchased from Origene (#RC203657, #RC235833, and #RC207956, respectively). To produce the Q2 domain of the Opi1 protein tagged with glutathione-S-transferase (GST), a cDNA encoding the Q2 domain of Opi1 (a gift from Dr. Alwin Köhler, University of Vienna, Austria) was cloned into pGEX 6P-1 (Amersham Pharmacia Biotech).

### Cell culture

H1299 cells, A549 cells, MCF7 cells, MB352 cells, MIA-PaCa-2 cells, and Panc-1 cells were obtained from the American Type Culture Collection, and cultured under 5% CO_2_ at 37 °C in Dulbecco’s modified Eagle’s medium supplemented with 10% fetal bovine serum. HMLE cells were generated by introducing SV40 large T-antigen and human telomerase reverse transcriptase into a primary culture of normal mammary epithelial cells (Lonza). HMLE cells were cultured in Mammary Epithelial Cell Growth Medium (Lonza). No mycoplasma were detected in cultures by 4′,6-diamidino-2-phenylindole (DAPI) or TO-PRO-3 (Thermo Fisher Scientific) staining. Polyclonal cell lines capable of the inducible expression of normal p53 (p53 WT) or a mutant p53 (p53 QM) were generated by infection of H1299 cells with the corresponding retrovirus and the rtTA retrovirus (Clontech), followed by selection with puromycin (1 µg/mL) and geneticin (G418, 500 µg/mL). The cells were cultured for 48 to 72 h in the presence of doxycycline (0.5 µg/mL) to induce the corresponding genes. Polyclonal cell lines expressing an epitope-tagged CTDNEP1 (MCF7-eCTDNEP1) were generated by the transfection of MCF7 cells with a vector encoding CTDNEP1, followed by selection with geneticin (G418, 250 µg/mL). H1299 and MB352 cells were stably transfected with pPB vectors encoding Fucci probes (Onodera et al., 2018).

### Antibodies and reagents

Primary antibodies for immunofluorescence, PLA, and immunoblot analysis were purchased from commercial sources, as listed in Supplementary file 3. Thymidine (Sigma, #T9250) was used to inhibit S-phase progression. DSP (Dojindo, #D629) was used to crosslink cellular proteins at 1 mM for 30 min at room temperature. Bis(sulfosuccinimidyl)suberate disodium salt (BS3; Dojindo, #B574) was used to crosslink the antibodies to protein G magnetic beads (Cell Signaling Technology). Benzonase (Merck, #70664-3CN) was added to the nuclear extracts at 100 units/mL with 1 mM MgCl_2_ overnight at 4 °C to digest nucleic acids. Deoxycytidine (Sigma, #D3897) was used to release cells from the thymidine-mediated blockade of cell cycle progression. MG132 (Enzo Life Sciences, #BML-PI102-0005) was used at 1 µM for 20 h to inhibit proteasomal function.

### Western blotting

Samples were loaded onto e-PAGEL gel (ATTO) or TGX FastCast gel (Bio-Rad), electrophoresed, and then the proteins in the gel were transferred onto Immobilon-FL polyvinylidene difluoride membranes (Millipore). The membranes were blocked with BlockPRO Protein-free Blocking Buffer (Visual Protein) or 1% bovine serum albumin (BSA) and 5% skim milk in phosphate-buffered saline (PBS), and then incubated with primary antibody solution at 4 °C overnight. After washing with PBS containing 0.1% Tween 20 (PBST), the membrane was incubated with IRDye secondary antibodies (LI-COR) or horseradish peroxidase-conjugated secondary antibodies (Jackson ImmunoResearch Laboratories) at room temperature for 1 h. To detect the immunoprecipitated endogenous CTDNEP1, TidyBlot^TM^ horseradish peroxidase-conjugated secondary antibody (BioRad) was used to enable the detection of immunoblotted target protein bands without interference from denatured IgG. After washing again with PBST, the membrane was scanned with Odyssey Imaging System (LI-COR) or detected with ImageQuant LAS 4000 mini imager (GE Healthcare), followed by its reaction with detection reagents (ECL Western Blotting Detection Reagents, GE Healthcare or SuperSignal^TM^ West Dura, Thermo Fisher Scientific). The primary antibodies used and their dilutions are listed in Supplementary file 3. All images were processed using Image J and/or Photoshop software (Adobe).

### Cell-cycle analysis

H1299 cells transduced with a vector encoding p53 WT or p53 QM were cultured with or without thymidine for 20 h. Cells were then trypsinized, washed twice in PBS, and fixed with ice-cold 70% ethanol for 2 h. Fixed cells were then washed twice in PBS, incubated with 0.5 mg/mL RNase A (Sigma, #R4642) at 37 °C for 30 min, followed by staining of DNA with 50 μg/mL propidium iodide. Stained cells were subjected to flow cytometric analysis using FACSVerse^TM^ (BD).

### Immunofluorescence analysis

For immunofluorescence analysis, cells cultured on coverslips were fixed either with 4% paraformaldehyde in PBS for 10 to 20 min followed by permeabilization with 0.1% Triton X-100 in PBS for 10 min, or fixed with 100% ethanol for 10 min at −20 °C, and incubated with blocking solution (1% BSA in PBS) for more than 60 min at room temperature. The cells were then incubated with primary antibodies for 60 min at room temperature, then washed with blocking solution and incubated with Alexa Fluor 488-, Alexa Fluor 647-, or Cy3-conjugated secondary antibodies (Molecular Probes) for 30 min. The cells were also stained with TO-PRO-3 dye or DAPI to visualize nuclei. The cells were finally washed with blocking solution, mounted onto glass slides with ProLong Diamond antifade reagent (Thermo Fisher Scientific), and observed using a confocal laser-scanning microscope with an oil-immersion objective (CFI Plan Apo VC 100×/1.4 NA or Apo λS 60×/1.4 NA), and analyzed with the attached software (Model A1 or A1R with NIS-Elements, Nikon), or with an N-SIM microscope (Nikon) with an oil-immersion objective (100×/1.49 NA), laser illumination, and an electron-multiplying charged-coupled device camera. Image reconstruction was performed using NIS-Elements software (Nikon). All images were processed using Photoshop software.

### Quantification of the perinuclear accumulation of H3K27me3

Each nuclear compartment was divided into three equally spaced concentric elliptic regions, starting from the boundary of the nucleus (selected manually using NIS-Elements software. The outermost concentric regions were taken as the perinuclear regions, and the innermost concentric regions were taken as the central nuclear regions. The average pixel intensities of H3K27me3 in these regions were measured by NIS-Elements software. The perinuclear accumulation of H3K27me3 was defined as their ratio (*i.e.*, the average H3K27me3 intensity of the perinuclear regions/the central nuclear regions) being more than 2. Nuclei with aberrant morphology (*i.e.*, nuclei to which ellipses do not fit) were omitted from this analysis (Figure 1–—figure supplement 1A).

### PLA

To visualize the subcellular localization of the H3K27me3-lamin A/C, H3K27me3-H4, H3.1/H3.2-lamin A/C, and H3-EZH2 interactions, H1299 cells with or without p53 QM expression were cultured with thymidine for 20 h to be arrested in the S phase. Cells were then fixed, permeabilized, and incubated with primary antibodies, as described in the immunofluorescence analysis section. PLA reactions were performed using Duolink^TM^ In Situ PLA Probe Anti-Mouse PLUS, Anti-Rabbit MINUS, and Detection Reagents Orange (Sigma), following the manufacturer’s instructions. After the ligation and amplification reactions, cells were further incubated with the Alexa Fluor 488-conjugated anti-lamin B1 antibody (Santa Cruz, #sc-365214AF488) and TO-PRO-3 dye to visualize the nuclear lamina and DNA, respectively. Specificities of the antibodies used for the PLA were confirmed by immunostaining performed at the same concentrations.

### Cell cycle synchronization and release

Cells were synchronized at the G1/S border or S phase by a single thymidine block (2 mM thymidine, 18–20 h) and then released by incubation in fresh media containing deoxycytidine (24 μM).

### Labeling of newly synthesized DNA

To label and visualize newly replicating DNA, the cells released from the thymidine block by incubation in fresh media containing deoxycytidine (24 μM) for 2 h were quickly rinsed with a hypotonic buffer (10 mM HEPES [pH 7.4] and 50 mM KCl), and incubated in this buffer containing 0.1 mM biotin-dUTP (Roche, #11093070910) for 10 min. Cells were then cultured in fresh culture medium for 20 min before fixation. Biotin-dUTP incorporated into the DNA replication sites were visualized by Alexa Fluor 488-conjugated streptavidin (Thermo Fisher Scientific, #S32354).

### Protein crosslinking and immunoprecipitation

For crosslinking, cells were washed with PBS and incubated with 1 mM DSP in PBS for 30 min at room temperature. Cells were then lysed with cell lysis buffer (10 mM HEPES [pH 7.4], 10 mM KCl, 0.05% NP-40) supplemented with a protease inhibitor cocktail (Merck, #4693116001), and incubated on ice for 20 min. The lysates were then centrifuged at 14,000 rpm for 10 min at 4 °C. The supernatants (cytosolic fraction) were removed and the pellets (nuclei) were resuspended in modified RIPA buffer (50 mM Tris [pH 7.4], 500 mM NaCl, 1 mM EDTA, 1% Triton X-100, 1% SDS, and 1% sodium deoxycholate) supplemented with the protease inhibitor cocktail, bath-sonicated, and incubated on ice for 20 min. The lysates were then centrifuged at 15,000 rpm for 10 min at 4 °C. The supernatants (crosslinked nuclear extracts) were diluted to yield 140 mM NaCl and 0.1% SDS solutions for immunoprecipitation. For immunoprecipitation, normal rabbit IgG or anti-H3 antibodies were crosslinked to protein G magnetic beads by BS3. The crosslinked nuclear extracts were then immunoprecipitated with antibody-conjugated beads overnight at 4 °C. Samples for MS analysis or western blot analysis were prepared by incubating and boiling the beads in SDS sample buffer.

### Identification of H3.1-interacting proteins by liquid chromatography-tandem MS (LC-MS/MS)

Samples were reduced with 10 mM TCEP at 100 °C for 10 min, alkylated with 50 mM iodoacetamide at an ambient temperature for 45 min, and then subjected to SDS-PAGE. Electrophoresis was stopped at a migration distance of 2 mm from the top edge of the separation gel. After CBB staining, protein bands were excised, destained, and cut finely prior to in-gel digestion with Trypsin/Lys-C Mix (Promega) at 37 °C for 12 h. The resulting peptides were extracted from gel fragments and analyzed using Orbitrap Fusion Lumos mass spectrometer (Thermo Scientific) combined with UltiMate 3000 RSLC nano-flow HPLC (Thermo Scientific). Peptides were enriched with μ-Precolumn (0.3 mm i.d. × 5 mm, 5 μm, Thermo Scientific) and separated on an AURORA column (0.075 mm i.d. × 250 mm, 1.6 μm, Ion Opticks Pty Ltd.) using the two-step gradient; 2% to 40% acetonitrile for 110 min, followed by 40% to 95% acetonitrile for 5 min in the presence of 0.1% formic acid. The analytical parameters of Orbitrap Fusion Lumos were set as follows: resolution of full scans = 50,000, scan range (m/z) = 350 to 1,500, maximum injection time of full scans = 50 msec, AGC target of full scans = 4 × 10^5^, dynamic exclusion duration = 30 sec, cycle time of data-dependent MS/MS acquisition = 2 sec, activation type = HCD, detector of MS/MS = ion trap, maximum injection time of MS/MS = 35 msec, AGC target of MS/MS = 1 × 10^4^. The MS/MS spectra were searched against a *Homo sapiens* protein sequence database (20,366 entries) in SwissProt using Proteome Discoverer 2.4 software (Thermo Scientific), in which peptide identification filters were set at a false discovery rate of less than 1%. Label-free relative quantification analysis for proteins was performed with the default parameters of Minora Feature Detector node, Feature Mapper node, and Precursor Ions Quantifier node in Proteome Discoverer 2.4 software.

### MMEJ-assisted gene knock-in using CRISPR-Cas9 with the precise integration into target chromosome (PITCh) system

For stable genomic integration of Dronpa with puromycin N-acetyltransferase or blasticidin deaminase into the carboxy terminus of H3.1, H1299 cells were transfected with two plasmids. The first plasmid encodes the inserted sequence flanked by two microhomology arms with a length of 40 bp and containing short PITCh sequences at their distal ends (modified based on pCRIS-PITChv2-FBL, Addgene #63672) (Sakuma et al., 2016). The second plasmid directs the expression of appropriate sgRNAs for cleaving the PITCh sequences and for targeting of the *HIST1H3A* locus to create a DNA fragment, and Cas9 (modified based on pX330A-1x2, Addgene #58766, and pX330S-2-PITCh, Addgene #63670) (Sakuma et al., 2016). From 3 days after transfection, the amount of puromycin (up to 0.7 μg/mL) or blasticidin (up to 7 μg/mL) was increased gradually for 20 days to enable the proliferation of cells with genomic insertion of the resistance gene.

### FRAP analysis of H3.1-Dronpa

To analyze nuclear recovery of the H3.1-Dronpa signals, their fluorescence in the nucleus was fully photobleached for 0.5 s using the 488-nm laser at its maximum (100%) intensity. Immediately after photobleaching, a 2- to 3-hour time series was acquired taking 1 image every 5 min. The intensity of the 488-nm laser was at 0.1% to 0.4% of the maximum, to minimize bleaching of the signal.

### Protein lipid overlay assay

Nitrocellulose membranes spotted with serial dilutions of different lipids (Membrane Lipid Arrays, Echelon Biosciences, #P-6003) were used to assess the lipid binding activity of histone H3.1. Membranes were incubated with blocking buffer (3% fatty acid-free BSA [Fujifilm, #017-15141] and 0.1% Tween 20 in PBS) for 1 h at room temperature before being incubated with 1.0 µg/mL of recombinant human histone H3.1 (New England Biolabs, #M2503S) or GST-fusion proteins for 1 h at room temperature. Alternatively, 1.0 µg/mL of recombinant human histone H3.1 was mixed with an equimolar amount of GST or recombinant human HSC70 (Abcam, #ab78431). The membrane was then washed with 0.1% Tween 20 in PBS, and incubated with an anti-histone H3.1/H3.2 antibody or anti-GST antibody for detection by immunoblotting.

### Phosphatase assay

To analyze the phosphorylation status of lipin 1, cells were lysed with 2× PPase reaction buffer (1× buffer contains 50 mM HEPES [pH 7.5], 100 mM NaCl, 10% glycerol, and 0.5% Triton X-100) supplemented with a protease inhibitor cocktail (Merck, #4693116001). The lysates were then bath-sonicated on ice for 5 min, and centrifuged at 14,000 rpm for 10 min at 4 °C. Total lysates (200 µg each) were incubated with or without 400 U lambda protein phosphatase (New England Biolabs, #P0753S) and 1 mM MnCl_2_ at 30 °C for 30 min. The reaction was stopped with 5× SDS sample buffer, and boiled for 5 min before western blot analysis.

### Quantification of PA in H1299 nuclei by LC-MS/MS

To prepare H1299 cell nuclei, cells were collected and washed with ice-cold PBS on ice. Cells were lysed with a hypotonic buffer (10 mM HEPES [pH 7.4], 1.5 mM MgCl_2_, and 10 mM KCl) supplemented with 1 mM DTT, a protease inhibitor cocktail (Merck, #4693116001), and a phosphatase inhibitor cocktail (Sigma, #04906837001), and then incubated on ice for 15 min. NP-40 at 0.5% (v/v) was then added to the lysates, the lysates were briefly vortexed, and centrifuged at 10,000 × *g* for 40 sec at 4 °C. The supernatant (cytosolic fraction) was removed and the pellet (nuclei) was resuspended in the hypotonic buffer (10 mM HEPES [pH 7.4], 1.5 mM MgCl_2_, and 10 mM KCl) supplemented with 1 mM DTT, the protease inhibitor cocktail, and the phosphatase inhibitor cocktail. The nuclei were then centrifuged at 10,000 × *g* for 40 sec at 4 °C, washed twice with ice-cold PBS, and then subjected to lipid extraction.

A 50 μL methanol solution containing 20 pmol each of C15:0/18:1 PA (Avanti Polar Lipids, #330721) and C12:0/13:0 PE (Avanti Polar Lipids, #LM1100) were added to each nuclear sample in 1.2 mL methanol in a glass tube, followed by the addition of ultrapure water (400 μL), 2 M HCl (400 μL), and 1 M NaCl (200 μL). After vigorous vortex-mixing, 2 mL CHCl_3_ was added, followed by further vortexing for 1 min. After centrifugation (1,200 × *g* for 3 min at room temperature), the lower organic phase (crude lipid extract) was collected and transferred to a new glass tube. Purified phospholipids were derivatized by methylation using a method described previously (Morioka et al., 2022). Briefly, 150 μL of 0.6 M (trimethylsilyl)diazomethane (Tokyo Chemical Industry, #T1146) was added to the purified phospholipid fraction prepared as above at room temperature. After 10 min, the reaction was quenched with 10 μL glacial acetic acid. The samples were mixed with 400 μL CHCl_3_ and 400 μL ultrapure water, followed by vortexing for 1 min. After centrifugation at 1,200 × g for 3 min, the lower phase was dried under a stream of nitrogen, and redissolved in 100 μL acetonitrile.

LC-MS/MS was performed using a triple quadruple mass spectrometer QTRAP6500 (ABSciex) and a Nexera X2 HPLC system (Shimadzu) combined with a PAL HTC-xt autosampler (CTC Analytics). The mass range of the instrument was set at 5 to 2,000 *m/z*. Spectra were recorded in the positive ion mode as [M+H]^+^ ions for the detection of PAs and PEs. The ion spray voltage was set at 5.5 kV, cone voltage at 30 V, and source block temperature at 100 °C. Curtain gas was set at 20 psi, collision gas at 9 psi, ion source gas pressures 1/2 at 50 psi, declustering potential at 100 V, entrance potential at 10 V, collision cell exit potential at 12 V, and collision energy values at 35 eV. Lipid samples (10 μL each) dissolved in acetonitrile were injected using the autosampler, and molecules were separated using a Bio C18 column (1.9 μm, 2.1 × 150 mm) at 60 °C. LC was operated at a flow rate of 100 μL/min with a gradient, as follows: 90% mobile phase A (methanol/acetonitrile/deionized water = 18/18/4 containing 5 mM ammonium acetate) and 10% mobile phase B (2-propanol containing 5 mM ammonium acetate) were maintained for 2 min, linearly increased to 82% mobile phase B over 17 min, and maintained at 85% mobile phase B for 4 min. The column was re-equilibrated to 10% mobile phase B for 10 min before the next injection. Data were analyzed by MultiQuant 3.0.2 software (ABSciex). Concentrations of each molecular species were calculated by dividing the peak area of each species with the peak area of the corresponding internal surrogate standard. The sum of each phospholipid molecular species was calculated, and the ratio (PA/PE) was presented in the Figure.

### RNA interference (RNAi) experiments

Silencer Select or Silencer Pre-designed siRNAs targeting coding sequences of human *TP53* mRNA (5′-GUAAUCUACUGGGACGGAATT-3′: p53 #1, and 5′-GGUGAACCUUAGUACCUAATT-3′: p53 #2), human *CTDNEP1* mRNA (5′-GUACCAAACUGUUCGAUAUTT-3′: CTDNEP1 #1, and 5′-GGAUCUGGAUGAGACACUUTT-3′: CTDNEP1 #2), human *LPIN1* mRNA (5′-GACUUUCCCUGUUCGGAUATT-3′: LPIN1 #1, and 5′-GGAGUGUCUUUGAAUAGAATT-3′: LPIN1 #2), human *TMEM255A* mRNA (5′-AAACCUUAUUGAGAACAAATT-3′: TMEM255A #1, and 5′-GGCUUUAAGGACAUGAACCTT-3′: TMEM255A #2), or Stealth siRNAs targeting coding sequences of human *EZH2* mRNA (5′-GACCACAGUGUUACCAGCAUUUGGA-3′: EZH2 #1, and 5′-GAGCAAAGCUUACACUCCUUUCAUA-3′: EZH2 #2) were obtained from Thermo Fisher Scientific. Silencer Select negative control siRNA and Stealth negative control siRNA were purchased from Thermo Fisher Scientific. For the transfection of siRNA, cells were plated at 15% to 20% confluence in 6-well plates together with 15 nM Silencer Select or Silencer Pre-designed siRNA and 3.5 µL of RNAi MAX (Thermo Fisher Scientific), or with 150 pmol of Stealth siRNA and 3.5 µL of RNAi MAX. They were then cultured in complete medium for 24 h before a second round of siRNA transfection.

### Quantitative reverse transcription (RT)-PCR analysis

Total RNA was extracted from cells using Trizol reagent (Thermo Fisher Scientific), purified with Direct-zol RNA Miniprep kit (Zymo Research), and aliquots (1 μg) of the RNA were subjected to RT with SuperScript IV polymerase (Thermo Fisher Scientific) or PrimeScript RTase (TaKaRa). TaqMan RT-PCR primers for *NEP1R1*, *TMEM255A*, *CTDNEP1*, and *ACTB* (encoding β-actin) were obtained from Applied Biosystems for quantitative PCR analysis. Quantitative PCR analysis was performed using the 7300 Fast Real-Time PCR System (Applied Biosystems). For the quantification of *NEP1R1*, *TMEM255A*, *CTDNEP1*, and *ACTB*, absolute quantification was performed using plasmids encoding human *NEP1R1*, *TMEM255A*, *CTDNEP1*, and *ACTB* as a standard, respectively.

### RNA-Seq for H1299 cells

Total RNA was isolated from each sample using Direct-zol RNA MiniPrep Kit (Zymo Research) together with DNase I treatment. The integrity and quantity of the total RNA was measured using an Agilent 2100 Bioanalyzer RNA 6000 Nano Kit (Agilent Technologies). Total RNA obtained from each sample was subjected to sequencing library construction using NEBNext Ultra Directional RNA Library Prep Kit for Illumina (New England Biolabs) with NEBNext Poly(A) mRNA Magnetic Isolation Module, according to the manufacturer’s protocol. The quality of the libraries was assessed using an Agilent 2100 Bioanalyzer High Sensitivity DNA Kit (Agilent Technologies). The pooled libraries of the samples were sequenced using NextSeq 500 (Illumina) in 76-base-pair single-end reads. Sequencing adaptors, low-quality reads, and bases were trimmed with Trimmomatic-0.32 tool (Bolger et al., 2014). The sequence reads were aligned to the human reference genome (hg19) using Tophat 2.1.1 (bowtie2-3.2.0) (Langmead and Salzberg, 2012), which can adequately align reads onto the location including splice sites in the genome sequence. Files of the gene model annotations and known transcripts, which are necessary for whole transcriptome alignment with Tophat, were downloaded from Illumina’s iGenomes website (http://support.illumina.com/sequencing/sequencing_software/igenome.html). The aligned reads were subjected to downstream analyses using StrandNGS 3.2 software (Agilent Technologies). The read counts allocated for each gene, and transcripts (RefSeq Genes 2015.10.05) were quantified using the Trimmed Mean of M-value method (Robinson and Oshlack, 2010).

### Statistical analysis

Statistical analysis was performed using GraphPad Prism 6 software (GraphPad Software, Inc.). Quantitative data are presented as the mean ± standard error of the mean. The sample size for each experiment and the replicate number of experiments are indicated in the Figures or Figure legends. Comparisons between two groups or more than two groups were performed using the Student *t*-test or one-way ANOVA followed by Dunnett’s multiple comparisons test, respectively. Statistical significance was indicated in the Figures or Figure legends. A *P*-value of less than 0.05 was considered to indicate a statistically significant difference between groups.

## Acknowledgments

We thank H.A. Popiel for her critical reading of the manuscript. We also thank Dr. Alwin Köhler and Dr. Anete Romanauska (University of Vienna, Austria) for providing the cDNA encoding the Q2 domain of Opi1. We would like to thank the Nikon Imaging Center at Hokkaido University for technical support.

This work was supported by JSPS KAKENHI grant numbers JP20K07305, JP20H05523, and JP16H06280 (to T.O.); JP18H02608 and JP21H02675 (to H.S.); JP20H03206 (to J.H.); JP20H04920 (to J.S.); JP20H03433 (to T.S.); SGH Cancer Research Grant (to T.O.); AMED grant 18gm0710002h0006 (to T.S.); and by Nanken-Kyoten (Grant No. 2023-kokunai 13 and 2024-kokunai 33), Tokyo Medical and Dental University (to T.O.).

## Author contributions

T.O. conceived the project and performed the imaging and biochemical experiments. A.H. performed the flow cytometric analysis. Y.O. generated the Fucci constructs and provided instruction on the PLA experiments. N.O. and K.U. performed the MS of the binding proteins of H3.1, and analyzed the data. J.H., J.S., and T.S. performed the MS of nuclear lipids, and analyzed the data. T.O. and H.H. performed the experiments of gene knock-in of the *HIST1H3A* locus. T.O. wrote the manuscript. H.S. supervised the project and modified the manuscript.

## Supplemental information

### Supplementary figure legends

**Figure supplement 1.**
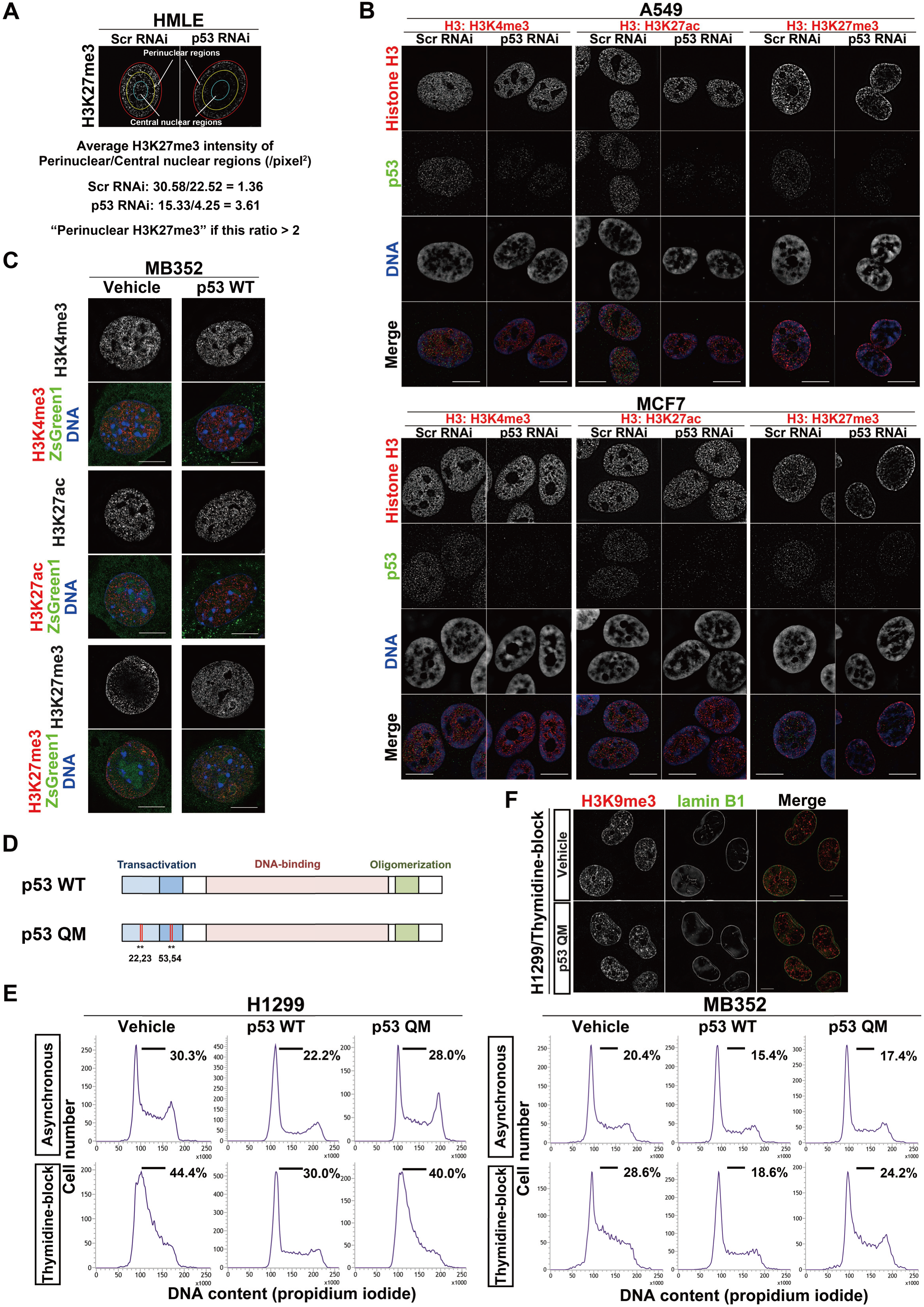
Loss of p53 induces perinuclear H3K27me3 accumulation, and p53 QM does not delay cell cycle progression in H1299 and MB352 cells. **A**, Definition of the perinuclear accumulation of H3K27me3 (see also Methods section). **B**, Representative SIM images showing the localization of H3K4me3, H3K27ac, or H3K27me3 (red), p53 (green), and DNA (blue), in A549 cells (top) and MCF7 cells (bottom). Bars, 10 µm. **C**, Representative SIM images of MB352 cells showing the localization of H3K4me3, H3K27ac, or H3K27me3 (red), and DNA (blue). ZsGreen1 (green) indicates expression of vehicle or p53 WT. Bars, 10 µm. **D**, Domain structure of p53. The human p53 QM used in this study harbors the L22Q/W23S/W53Q/F54S mutations (indicated as *) in the transactivation domain. **E**, Representative flow cytometry histograms of H1299 cells (left) and MB352 cells (right) are shown. Cells in the G1/S phase are defined as those under the black horizontal bar, and their proportions are indicated. **F**, Representative SIM images of H1299 cells showing the localization of H3K9me3 (red) and lamin B1 (green). Bars, 10 µm

**Figure supplement 2.**
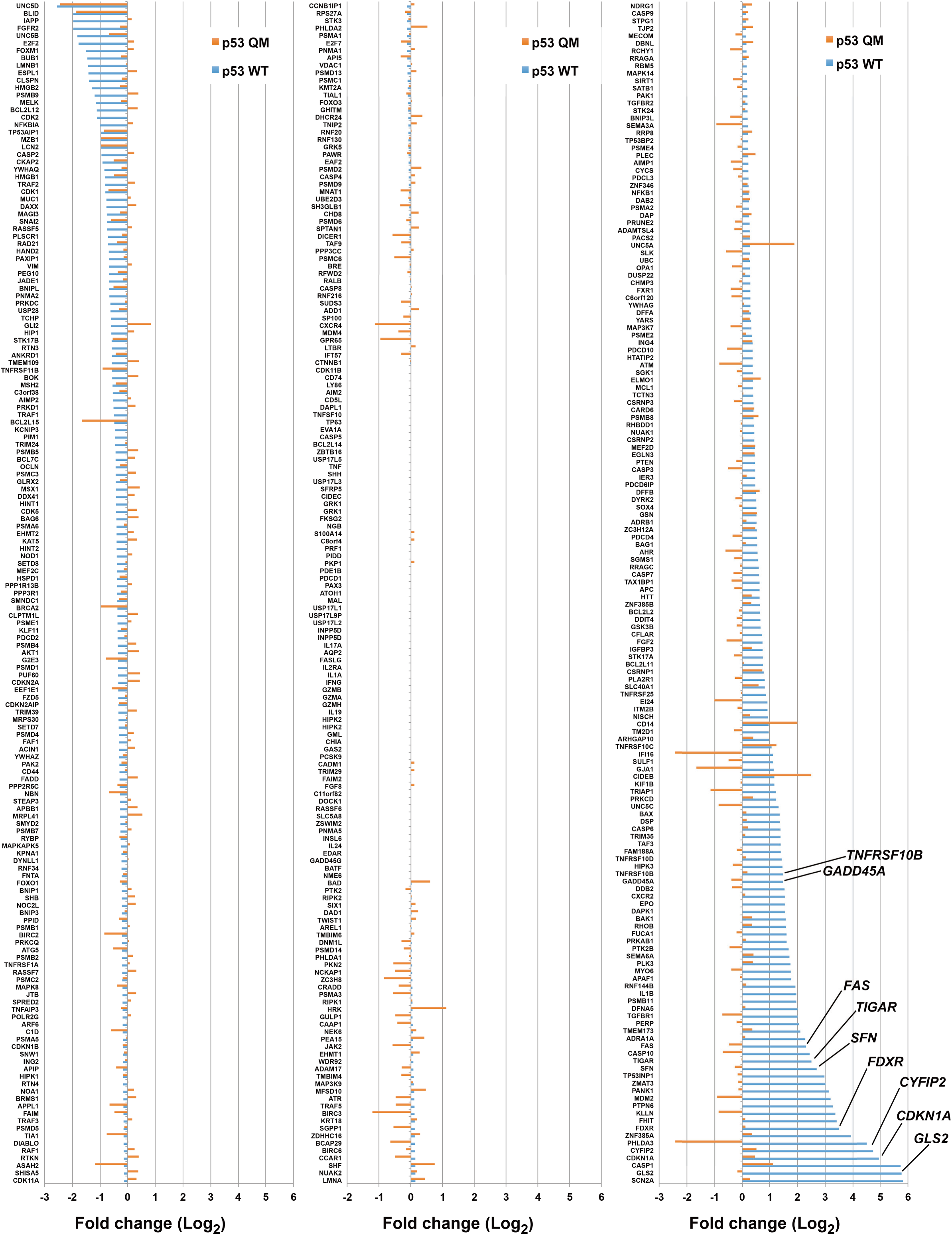
Changes in expression levels of p53-target genes upon p53 WT or p53 QM expression in H1299 cells. Genes identified by RNA-Seq analysis are listed according to the Gene Ontology term that includes “p53”. Some other genes are also listed from reports in the literature (Fischer, 2017). The fold changes (Log_2_) to the control are indicated. The expression of p53 WT but not p53 QM induces a wide range of known p53-target genes, such as those implicated in cell cycle arrest (e.g., *CDKN1A* encoding p21/Cip1 (el-Deiry et al., 1993), *SFN* encoding 14-3-3 sigma (Hermeking et al., 1997), and *GADD45A* (Kastan et al., 1992)), apoptosis (e.g., *CYFIP2* (Jackson et al., 2007), *FAS* (Muller et al., 1998), and *TNFRSF10B* encoding DR5 (Wu et al., 1997)), and metabolism (e.g., *GLS2* (Hu et al., 2010; Suzuki et al., 2010), *FDXR* (Liu and Chen, 2002), and *TIGAR* (Bensaad et al., 2006)).

**Figure supplement 3.**
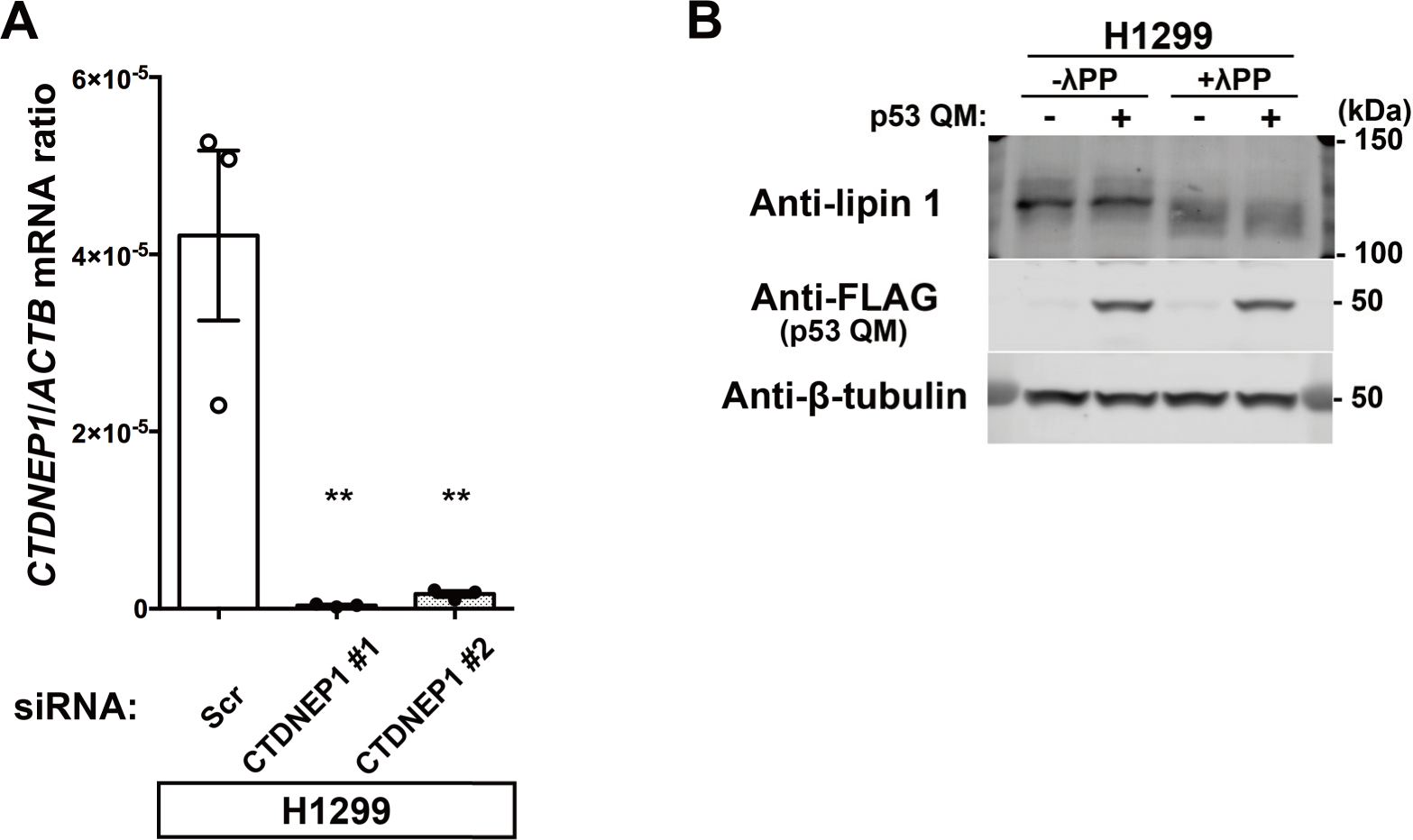
*CTDNEP1* silencing and phosphorylation status of lipin 1 in H1299 cells. **A**, Quantification of *CTDNEP1* mRNA (normalized by *ACTB* mRNA) in H1299 cells. n = 3 biological replicates; \*\**P* < 0.01; one-way ANOVA followed by Dunnett’s multiple comparisons test. **B**, Representative immunoblots using the indicated antibodies. λPP, lambda protein phosphatase.

**Figure supplement 4.**
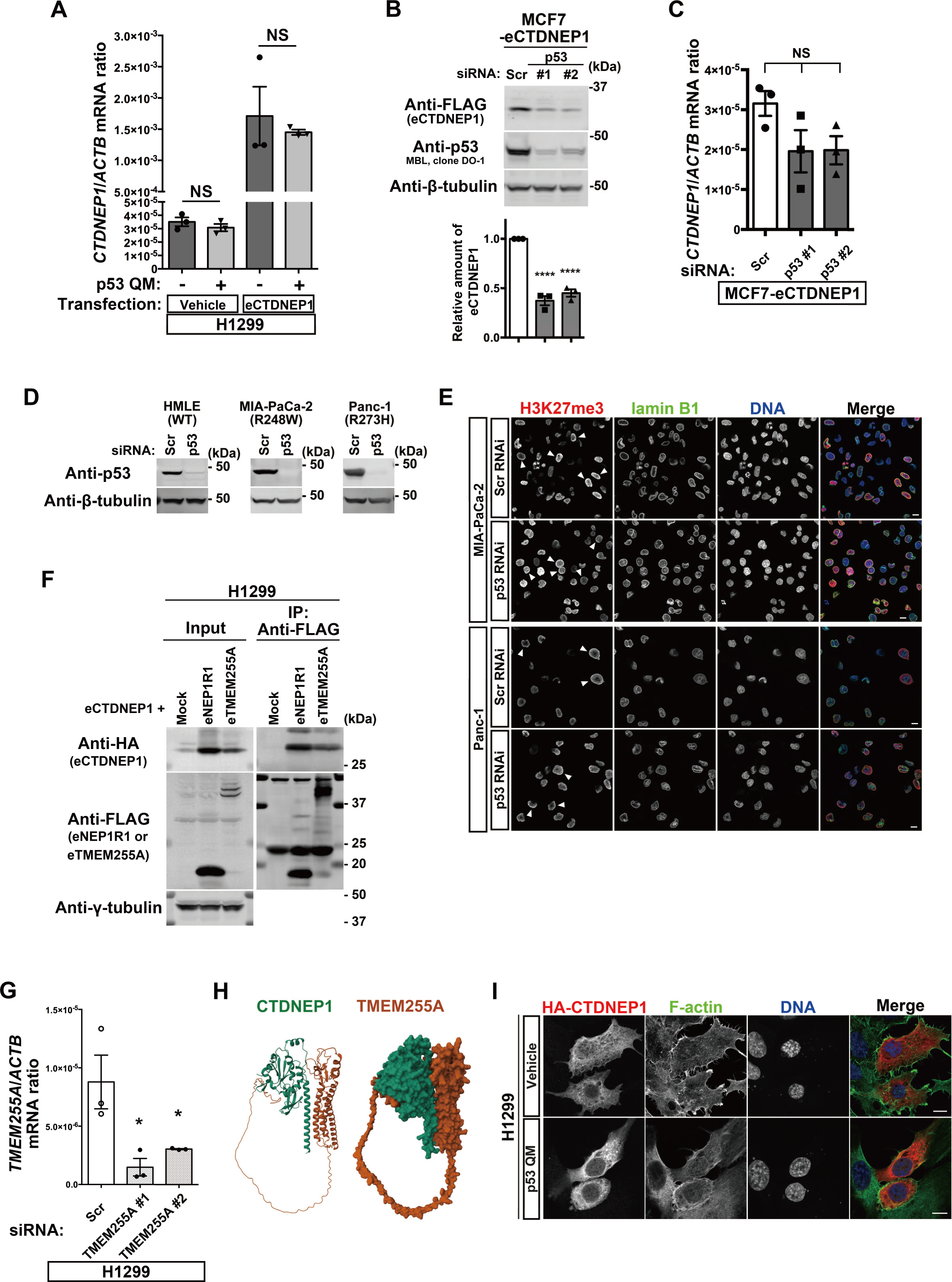
p53 increases CTDNEP1 via the upregulation of *TMEM255A*. **A**, Quantification of *CTDNEP1* mRNA (normalized by *ACTB* mRNA) in H1299 cells. N = 3 biological replicates; NS, not significant; two-tailed, unpaired *t*-test. **B**, Representative immunoblots using the indicated antibodies. Normalized eCTDNEP1/β-tubulin or eCTDNEP1/GAPDH ratios are shown. n = 3 biological replicates; \*\*\*\**P* < 0.0001; one-way ANOVA followed by Dunnett’s multiple comparisons test. **C**, Quantification of *CTDNEP1* mRNA (normalized by *ACTB* mRNA) in MCF-7 cells. n = 3 biological replicates; NS, not significant; one-way ANOVA. **D**, Representative immunoblots using the indicated antibodies. **E**, Representative confocal images of MIA-PaCa-2 and Panc-1 cells showing the localization of H3K27me3 (red), lamin B1 (green), and DNA (blue). Bars, 10 µm. Arrowheads indicate the cells with perinuclear H3K27me3. **F**, Representative immunoblots using the indicated antibodies. IP, immunoprecipitation. **G**, Quantification of *TMEM255A* mRNA (normalized by *ACTB* mRNA) in H1299 cells. n = 3 biological replicates; \**P* < 0.05; one-way ANOVA followed by Dunnett’s multiple comparisons test. **H**, Structure of the CTDNEP1-TMEM255A complex predicted by ColabFold. A molecular ribbon representation (left) and a molecular surface plot (right) are shown. **I**, Representative confocal images of H1299 cells showing the localization of HA-CTDNEP1 (red), F-actin (green), and DNA (blue). Bars, 10 µm.

### Video legends

**Video 1. FRAP analysis of H3.1-Dronpa in the absence of p53 in H1299 cells**

H1299 cells harboring the *Dronpa* gene at the *HIST1H3A* locus were thymidine-blocked and photobleached (frames 1, 2, and 3). Images were then taken every 5 min for 2 to 3 h. The frame interval is 300 msec (1,000× speed).

**Video 2. FRAP analysis of H3.1-Dronpa in the presence of p53 QM in H1299 cells**

H1299 cells harboring the *Dronpa* gene at the *HIST1H3A* locus and expressing p53 QM were thymidine-blocked and photobleached (frames 1, 2, and 3). Images were then taken every 5 min for 2 to 3 h. The frame interval is 300 msec (1,000× speed).

**Supplementary file legends**

**Supplementary file 1. List of genes upregulated or downregulated upon the expression of p53 WT or p53 QM in H1299 cells**

**Supplementary file 2. H3.1 interacting proteins identified by MS analysis**

**Supplementary file 3. List of antibodies used in this study**

